# Identifying Factors Important for Conservation at Sites of Synonymous Variations

**DOI:** 10.1101/2024.01.01.573819

**Authors:** Abhirami Ramasubramanian, Uma Sunderam, Rajgopal Srinivasan

## Abstract

Synonymous mutations can have a deleterious effect leading to disease, even though they are not protein altering. Variations at genomic sites leading to synonymous variants are frequently highly conserved across species. Several prediction methods have been developed to assess the impact of synonymous mutations and are highly dependent on having validated sets of both deleterious and benign synonymous mutations. However, validated data available for deleterious synonymous mutations is sparse unlike for missense mutations. Rather than develop a model for predicting pathogenicity of synonymous variants, we seek to understand the relative importance of various factors that lead to conservation at sites of synonymous variants. Our study built machine learning models using various features on a large set of reported and generated synonymous variants (Zeng Z et al, 2019) to predict conservation (Genomic Evolutionary Rate Profiling – Rejected Substitution (GERP RS) base scores and Phylogenetic p-values for 100 vertebrates (PP100)) at genomic sites. We used the extreme gradient boosting classifier to classify sites as high, medium and low conservation at different cutoffs. Our experiments report an AUC between 0.74-0.79 and the sensitivity was significant. Of the features we explored, a few alternate allele independent properties were repeatedly flagged as having high impact. These findings provide information for predictors to further improve models for synonymous variant impact.

## Introduction

Variants causing a direct change in the transcribed protein such as missense, nonsense, frame-shift insertion deletions, and splice site disrupting constitute the majority of pathogenic variants in many disease databases. Synonymous mutations can also have a deleterious effect on function even though they are not protein altering. Variations at synonymous sites can impact splicing including cryptic splice, splice enhancers and suppressors, disrupt transcription, co-translational folding and mRNA stability among other things. Several machine learned models have been developed to assess the impact of synonymous mutations and are highly dependent on having validated sets of deleterious and benign synonymous mutations (Zeng Z et al, 2019; Buske OJ et al, 2013; Livingstone M et al, 2017; Livingstone M et al, 2017; Shi F et al, 2019). However, validated data available for deleterious synonymous mutations is sparse unlike for missense mutations. For e.g. ClinVar database (accessed 30^th^ April, 2022) reports only 318 pathogenic synonymous variants in comparison to 56,100 pathogenic non-synonymous variants (missense and nonsense).

*Rather than develop a model for predicting pathogenicity of synonymous variants, we seek to understand the relative importance of various factors that lead to conservation at sites of synonymous variants.* Our study built machine learning models using a variety of features on a large set of reported and generated synonymous variants that were used in another study (Zeng Z et al, 2019) to predict the conservation score at the variant site. As evidenced (fig 1a) by analyzing synonymous ClinVar mutations, GERP RS (Rejected Substitution) base scores (Cooper GM et al, 2005; Davydov EV et al, 2010) and PhyloP values for 100 vertebrates (PP100) (Pollard, K.S. et al, 2009) provide good, albeit imperfect, discrimination between benign and pathogenic variants. We therefore postulate that factors important for predicting conservation may also play an important role in predicting functional deleteriousness. We report the results of our study.

**Figure 1:**
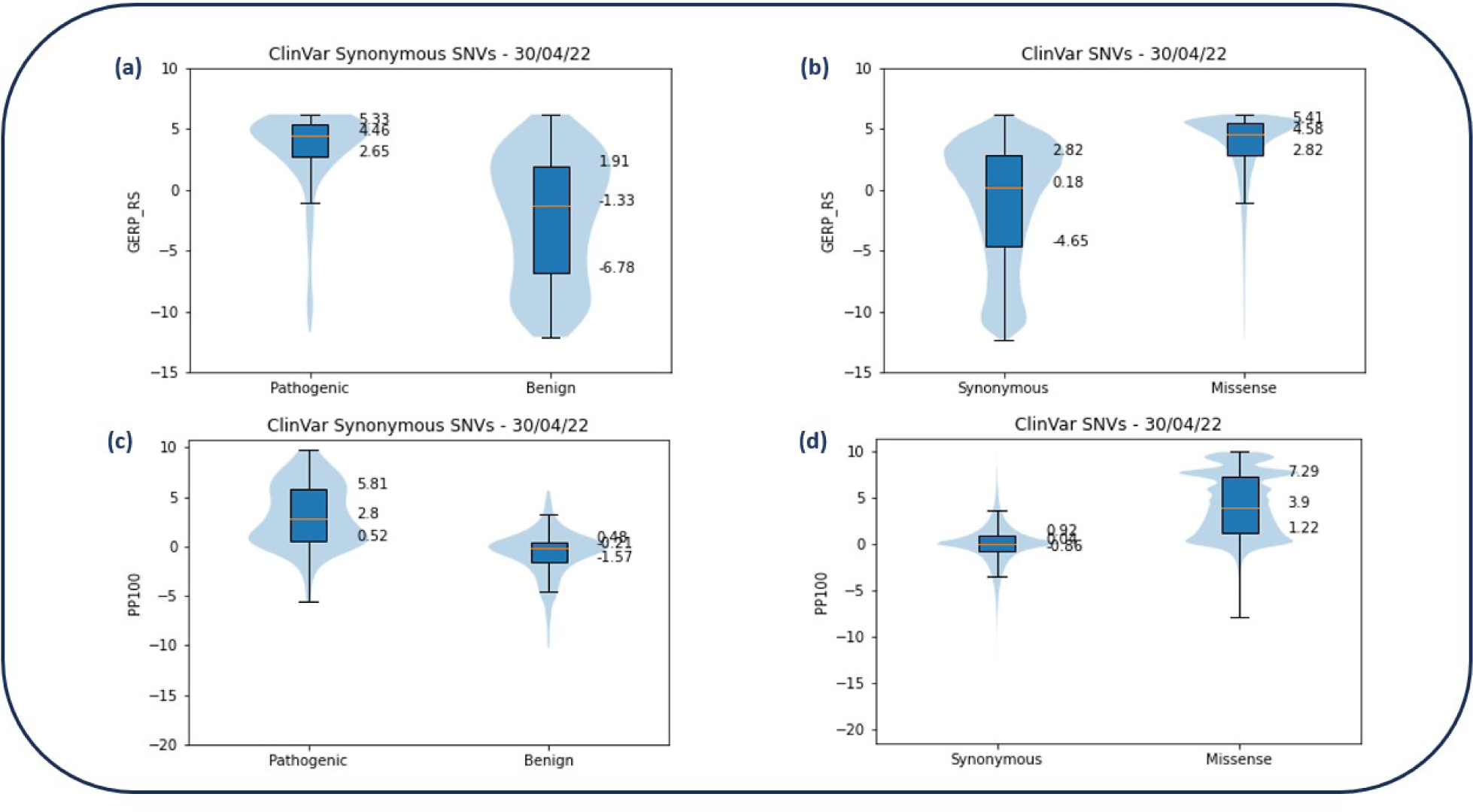
**a)** GERP RS distributions of benign and pathogenic ClinVar synonymous variants **b)** GERP RS distributions of synonymous and missense ClinVar variants **c)** PP100 distributions of benign and pathogenic ClinVar synonymous variants **d)** PP100 distributions of synonymous and missense ClinVar variants

## Materials and Methods

### Datasets

#### (i) Synonymous Variants data

Observed and generated synonymous variants that were used in a previous study (Zeng Z et al, 2019) were combined to form the data set for the study. The observed variants (1,362,607 synonymous variants) were collected from public databases such as 1000 Genomes (The 1000 Genomes Project Consortium, 2015), ExAC (Lek M et al, 2016) and gnomAD (Karczewski, K.J. et al 2020). The generated variants were a size matched random subset of all possible synonymous variants. Duplicates in the combined data were excluded.

#### (ii) Variant Selection

A genomic variant site can have a different mutation type depending on the transcript. We selected our synonymous variants from the above set considering only those synonymous variants that occured in Matched Annotation from NCBI and EMBL-EBI (MANE) transcripts (Morales J. et al, 2022). The annotations were from GENCODE (Frankish A et al, 2021) project (version 39 lifted over to GRCh37) and mutation effect as annotated from our in-house script Varant (http://compbio.berkeley.edu/proj/varant/Home.html).

In order to restrict the variant effect to purely synonymous sites, we only retained those where all three possible allele changes were synonymous in the thus prioritized transcript.

Variant sites that were within 3 bases of splice acceptor or donor sites were excluded from our experiments to avoid any bias caused by direct disruption of the splice region. Moreover, only variants that had annotations for all the features were retained. For e.g. if by virtue of being in the first or last exon, a variant did not have a splice acceptor or donor score for the exon, it was removed from the data set. The final dataset consisted of 7,24,568 sites of synonymous variation.

### Features

A total of 35 features were used and briefly described below.

#### (i) Position based

Features that describe the location of the site include: 1) the position of the site in the exon, 2) the position of the corresponding amino acid in the protein, and 3) whether the site falls in functional regions such as transcription factor binding sites.

#### (ii) Splice based

Many variants have deleterious effects because of their impact on splicing. Some of the splicing-related features we have considered include: 1) the MaxEntScan (Yeo G et al, 2004; Eng L et al, 2004) donor and acceptor score percentiles of the splice sites and 2) whether or not cryptic splice sites were created by any of the alternate alleles at that position and if they were, their respective MaxEntScan scores. The MaxEntScan percentiles were calculated by computing the MaxEntScan scores of all unique splice acceptors and donors from Gencode transcripts.

We also considered the impact of the variants on Exonic Splice Enhancers (ESE) and Exonic Splice Silencers (ESS) (Fairbrother WG et al, 2002; Wang, Z et al, 2004). A count of the number of ESEs lost and ESSs gained and the ESR scores (average hexamer ESRseq scores (Ke S et al, 2011) computed over a sliding window over the site) for the reference and alternate alleles as well as their difference were the features used to study this.

#### (iii) Codon based

The codon based features consist of codon usage features such as the Relative Synonymous Codon Usage (RSCU) calculated for the reference codon and all alternate codons, along with their respective differences. RSCU is defined as the ratio of the observed frequency of codons to the expected frequency given that all the synonymous codons for the same amino acids are used equally (Eq 1) (Nakamura Y et al, 2000). We have calculated RSCU globally as well as specific to the transcript considered for each site.

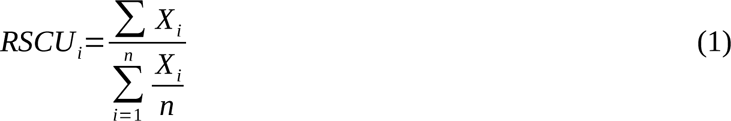

where Xi is the number of occurrences and RSCUi is the relative synonymous codon usage for codon i.

Codon related features that could potentially impact translation were also added to the set. Specifically, the additional occurence of the reference codon before or after and contiguous occurence with respect to the variant codon site.

To assess the impact of the actual codon, the reference codon was also encoded base-wise in terms of the purine/pyrimidine bases and double or triple hydrogen bondedness.

All the features are described in detail in Supplementary Table 1.

**Table 1:** Average metrics from 10-fold cross validations and 1 run of the independent test sets for the 4 different models.

| Dataset | Cutoff for high class | AUC | Precision | Recall | F1-score |
| --- | --- | --- | --- | --- | --- |
| Training data | GERP RS ≥ 90 percentile | 0.756 | 0.440 | 0.565 | 0.439 |
| | GERP RS ≥ 75 percentile | 0.741 | 0.525 | 0.559 | 0.529 |
| | PP100 ≥ 90 percentile | 0.786 | 0.467 | 0.600 | 0.480 |
| | PP100 ≥ 75 percentile | 0.742 | 0.529 | 0.558 | 0.535 |
| Test data | GERP RS ≥ 90 percentile | 0.757 | 0.438 | 0.568 | 0.434 |
| | GERP RS ≥ 75 percentile | 0.744 | 0.525 | 0.561 | 0.529 |
| | PP100 ≥ 90 percentile | 0.790 | 0.467 | 0.606 | 0.478 |
| | PP100 ≥ 75 percentile | 0.745 | 0.530 | 0.561 | 0.537 |

### Alternate allele features

Each sample represented a site rather than a variant since the conservation score was for the site and not alternate allele dependent. Of the alternate allele specific features, the variant which had the hypothesised maximum impact, for e.g., the maximum ESS gain among 3 alternate alleles was selected for each site.

Similarly, minimum of the ESR and RSCU alternate allele scores were considered amongst the variants. Variants creating cryptic donor sites were considered as having higher impact over those creating cryptic acceptor sites. Within cryptic splice sites of the same type, the allele that resulted in the maximum maxent cryptic donor or acceptor score percentile was selected.

### Model Building

Models were built using two different measures of conservation as the target variable. In one, the base-wise GERP RS conservation scores were used as the target variable and in another, PhyloP values for 100 vertebrates (Pollard, K.S. et al, 2009). Based on the observed distribution (Figures 1b and 1d) of reported synonymous variants in clinvar database, the data was divided into 3 classes of variants based on their evolutionary conservation scores. These corresponded to the high class (highly conserved), low class (low conservation) and medium class (variants with conservation scores between the high and low classes). Two different distributions of the data were used: Case 1) high class having conservation scores above the 90th percentile and the low class with conservations scores in the bottom 10^th^ percentile and Case 2) high class scores above the 3^rd^ quartile and low class conservation scores below the 1^st^ quartile (cutoffs in Supplementary Table 2). The remaining variants in between the high and low classes were considered the medium class in both cases. These precomputed scores were obtained from http://mendel.stanford.edu/SidowLab/ <u>downloads/gerp/hg19.GERP_scores.tar.gz</u>in the case of GERP RS and https://hgdownload.soe.ucsc.edu/goldenPath/hg19/phyloP100way/hg19.100way.phyloP100way.bw in case of PhyloP100 way. Regression models were also built and compared (results not shown) but were discarded in favour of the multiclass classifier because of inadequate prediction accuracy.

**Table 2:** Significant features reported by SHAP and feature importances.

| Feature group | Features |
| --- | --- |
| Codon usage related | rscu_ref, rscu_tx_ref, rscu_tx_diff_min, rscu_diff_min, rscu_alt_min, rscu_tx_alt_min |
| Splice related | ESRREF, ESRDIFF_min, ESRALT_min, ese_loss_max, ess_gain_max, dist_5_prime, dist_3_prime |
| Reference codon related | rcodon2b, rcodon3b, rcodon3p, rcodon2p |
| Others | TFBS, exon_length, codon_count |

The XGBoost package (Chen T et al, 2016) version 1.7.3, a gradient boosting library designed to be a highly efficient implementation, was used to train and test the data. 3 class models based on the above data were built with number of estimators set to 1000 and other default parameter values.

## Results

### Model Evaluation

The model was evaluated with both 10-fold stratified cross validation as well as an independent test set (a randomly selected 20% of the total dataset). The model distinguished highly conserved and non-conserved variants with an AUC of between 0.74 and 0.79 (Table 1).

Similar AUC and balanced accuracy values were observed for the test set (Table 1).

### Significant Features

We next looked into significant features and their relative importances for the 4 models with SHapley Additive exPlanations (SHAP) values (Lundberg, S.M. et al, 2020), an approach to explain the output of machine learning models based on classical Shapley values from cooperative game theory and their related extensions. For each model, the top 15 features across the 3 classes were considered and a union across the 4 models resulted in 17 significant features. The top 15 features across the 3 classes in one of the models is shown in Figure 2. In addition, the top 15 features from feature importances module in *scikit learn* were also compared from the 10-fold cross validation data and 3 additional exclusive features were added to the list. The significant features are consolidated in Table 2.

**Fig 2:**
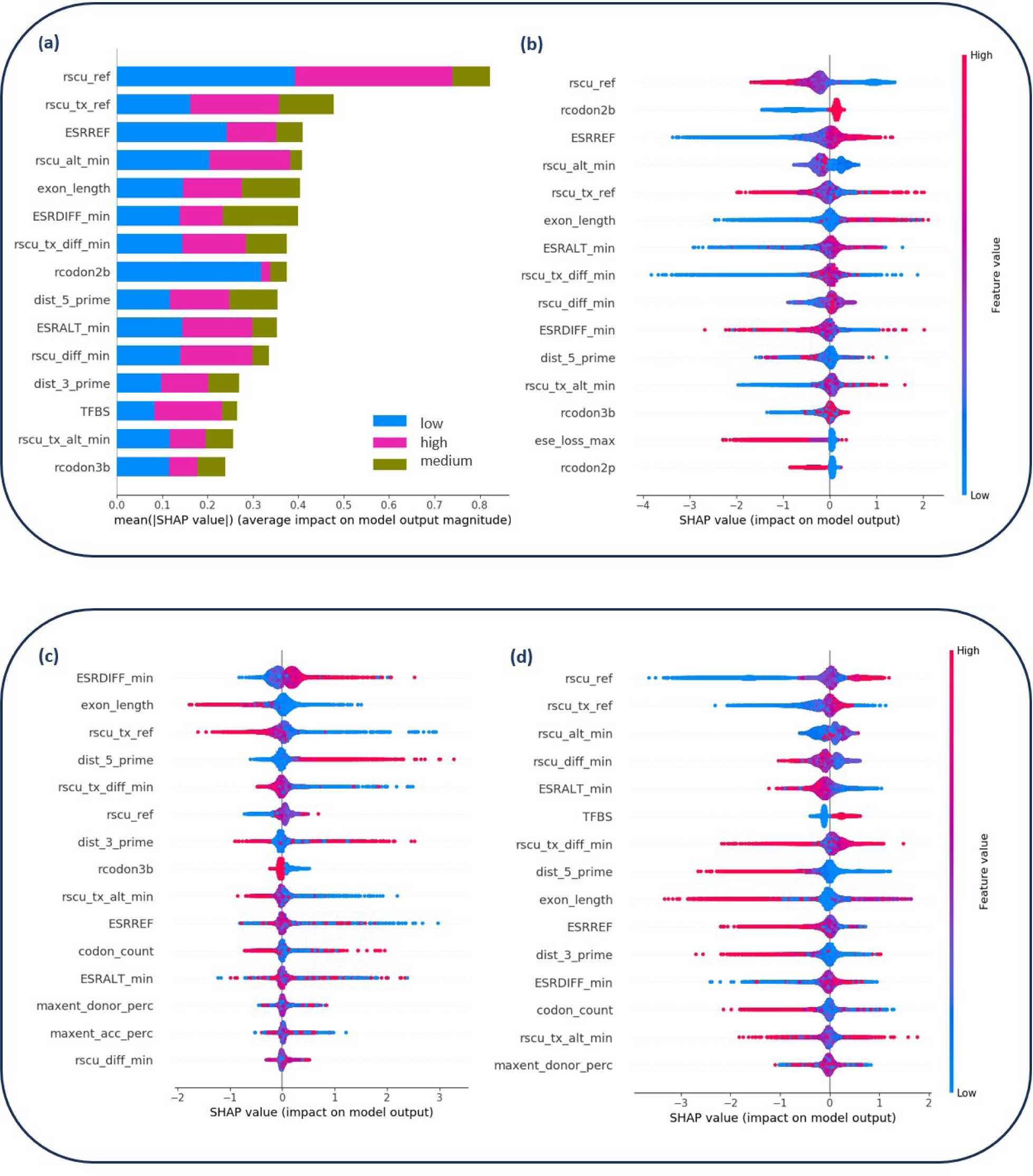
Shap Value plots for PP100 target model with >= 90 percentile high class. Important features based on mean SHAP values across the 3 different classes (box plot) and individual classes (beeswarm plots). clockwise from top left: **a)** box plot for significant features **b)** beeswarm plot for high class **c)** beeswarm plot for low class and **d)** beeswarm plot of middle class

The features that were consistently reported as significant were related to RSCU scores (both global and transcript based), ESR scores, base identity at substitution site, TFBS, exon length and codon count. RSCU (global value) for the reference allele was the top feature in 3 out of 4 models as per feature importances. It was also observed that for other feature groups like RSCU (transcript based) and ESR score, the reference feature frequently ranked highest among the set, ahead of the difference with the alternate allele and alternate allele values. The identity of the 2^nd^ and 3^rd^ reference codon base were also consistently in the top 15 features. Other features that were reported as significant albeit lower ranked in the top 15 were TFBS, exon length and codon count. The models were also built based on annotations using GRCh38 reference databases and similar trends were observed (Supplementary Table 3).

**Table 3:** Significant related feature groups performance on the independent test set.

| Features | AUC | Precision | Recall | F1-score |
| --- | --- | --- | --- | --- |
| Codon usage related group | 0.750 | 0.429 | 0.562 | 0.401 |
| Splice related group | 0.682 | 0.409 | 0.500 | 0.390 |
| Reference codon related group | 0.718 | 0.405 | 0.524 | 0.318 |
| All 3 groups together | 0.785 | 0.460 | 0.604 | 0.465 |
| Top 10 features (single feature models) | 0.778 | 0.453 | 0.597 | 0.453 |
| Top 20 features | 0.789 | 0.464 | 0.607 | 0.473 |

We next examined how groups of related features and individual features performed for a model built with PP100 as target and high class >= 90 percentile and low class < 10 percentile. We also built models with all the 20 short listed features and with only the top 10 best performing features based on the individual feature models. The results of the test run are reported in Table 3.

Of the 3 related feature groups, Codon usage performed the best with an AUC of almost 0.75. A few individual features – rscu_ref and rscu_diff_min marginally outperformed both the reference codon related and splice related groups (Supplementary Table 4). Reducing the feature set to only the top 10 features based on the single feature models (not shown), gave a performance almost on par with the full feature set. With all 3 related feature groups together (17 features), the performance difference with using all 35 features was negligible.

**Table 4:** Test set results for Models built for single exon, first exon and last exons.

| Model | Cutoff for high class | AUC | Precision | Recall | F1-score |
| --- | --- | --- | --- | --- | --- |
| Single exon models | GERP RS $\geq 75$ | 0.699 | 0.500 | 0.510 | 0.504 |
| | GERP RS $\geq 90$ | 0.736 | 0.475 | 0.484 | 0.479 |
| | PP100 $\geq 75$ | 0.712 | 0.507 | 0.519 | 0.512 |
| | PP100 $\geq 90$ | 0.741 | 0.475 | 0.493 | 0.483 |
| Last exon models | GERP RS $\geq 75$ | 0.739 | 0.535 | 0.556 | 0.541 |
| | GERP RS $\geq 90$ | 0.767 | 0.477 | 0.534 | 0.496 |
| | PP100 $\geq 75$ | 0.738 | 0.529 | 0.551 | 0.537 |
| | PP100 $\geq 90$ | 0.767 | 0.480 | 0.531 | 0.498 |
| First exon models | GERP RS $\geq 75$ | 0.697 | 0.490 | 0.506 | 0.495 |
| | GERP RS $\geq 90$ | 0.723 | 0.460 | 0.486 | 0.470 |
| | PP100 $\geq 75$ | 0.696 | 0.489 | 0.503 | 0.494 |
| | PP100 $\geq 90$ | 0.737 | 0.473 | 0.498 | 0.484 |

### Region-wise models

In order to minimize features without data, our model had excluded variants located in the first exon, last exon, and variants located in single exon genes. In order to include such variants and verify the model capability of distinguishing high, medium and low conservation, we built specific models for such regions with only applicable features.

Models for all 3 special cases of variants; viz; occuring in the first, last and single exon genes could distinguish between the high, medium and low conservation (Table 4). The model for first exon performed slightly below par with an AUC of between 0.69 to 0.74 compared to the other models. The last exon model could pick higher conservation variants better compared to all other models (precision).

Next, top 10 significant features that impacted the prediction were evaluated using SHAP as before. Although the overlap with features identified by the general models were similar, the order of importance of few individual features were different (Supplementary Figure 1). For e.g length of exon rose in significance especially for the single exon models and position in the exon also had a higher weightage.

### Significant Features with Other Predictors

In our experiments thus far, we had consistently observed some features as significant. These included Codon usage related, ESR scores and actual nucleotide at the third position of the codon. To assess the impact of a subset of features on available predictor algorithms, we experimented with adding combinations of our features to other features used by the predictors and also replacing the conservation component with the features to evaluate the performance.

The Identification of Deleterious Synonymous Variants (IDSV) model (Shi F, et al 2019) uses the random forest method to identify deleterious synonymous variants with 10 optimized features. We have attempted replicating their results with their data and code and further annotated their datasets with significant features identified in this study. We test whether top features can replace their conservation feature, PhyloP, and still retain performance. Since our main model excluded near splice variants, the variants selected for the training and test sets from the IDSV data excluded such variants. We retained synonymous SNVs from the IDSV train and test sets which occured in MANE transcripts. This resulted in a total of 418 variants in the training set (down from their original training set of 600), with 163 deleterious and 255 benign variants. Since we couldn’t replicate their Translation Efficiency feature values, we calculated these values using the MANE transcript that was selected for the variant. All other features were annotated and models were run as specified in their README and using the downloaded scripts. This experiment was carried out with the variant information and not the genomic sites as with our models since IDSV is a variant effect predictor.

Our test data for the experiment comprised of the IDSV test set, augmented with an independent set of pathogenic and benign synonymous ClinVar variants (accessed 30th April 2022). The synonymous clinvar variants were shortlisted as follows:

‘Pathogenic’ & ‘Pathogenic/Likely_pathogenic’ sSNVs for the deleterious class and ‘Benign’ sSNVs for the benign class were selected from the file, without checking for assertion. The ClinVar variants were filtered in the same way as the IDSV train and test sets as described previously and then annotated with the 10 IDSV features as well as our top 19 features (excluding TFBS as IDSV uses it as one of its 10 optimized features). The unique set of combined IDSV test and ClinVar variants not found in the train set was the test set used for our experiment. The combined test set had 163 deleterious and 39,448 benign sSNVs.

We observed that all metrics reported were lower when PhyloP was dropped as a feature (Table 5). However when the individual features shortlisted in our model were added, for most of the cases, the AUC increased (Supplementary Table 5). The performance after adding the 3 related features groups individually was comparable to the original dataset with PhyloP. Next, the top 10 best performing features, 3 related feature groups combined, and all shortlisted 20 features were added and the prediction run for the test set. In these cases, the performance exceeded that of the original model with all 10 IDSV features (including PhyloP).

**Table 5:** IDSV performance (independent test set combining IDSV test and clinvar variants) with different combinations of feature groups identified as significant in place of conservation score PhyloP 100 way conservation score feature (average values over 10 runs)

| **Dataset** | **AUC** | **Recall** | **Specificity** | **Precision** | **F1-score** |
| --- | --- | --- | --- | --- | --- |
| IDSV features including PhyloP | 0.716 | 0.510 | 0.791 | 0.009 | 0.017 |
| **Feature Groups added in place of PhyloP** |  |  |  |  |  |
| None (Baseline) | 0.699 | 0.499 | 0.784 | 0.008 | 0.016 |
| rscu related group | 0.717 | 0.499 | 0.801 | 0.009 | 0.017 |
| splice related group | 0.719 | 0.463 | 0.799 | 0.008 | 0.016 |
| rcodon related group | 0.703 | 0.485 | 0.797 | 0.008 | 0.016 |
| Top 10 features (from single feature models) | 0.730 | 0.514 | 0.825 | 0.010 | 0.020 |
| All 3 groups together | 0.736 | 0.543 | 0.814 | 0.010 | 0.020 |
| Top 20 feat | 0.734 | 0.524 | 0.817 | 0.010 | 0.020 |

## Discussion

We evaluated conservation classification as a proxy for variant deleterious effect using 35 features categorized into codon related, potential splicing disrupting and relative position within the protein. A large dataset with variants that were a mixture of reported and possible synonymous mutations were used for the multi class (high, medium and low conservation) classification and showed AUC, precision, recall and F1-score values that were higher than with random classification. Both 10-fold stratified cross validation and evaluation with an independent test set gave similar results.

When importances of the features in the model were examined, the top features were consistent across cutoffs and over multiple runs of the cross validation. It was notable that some significant features were reference based and ranked as important along with the alternate allele value and the difference between the reference and alternate allele of the related feature, the most striking being RSCU. Codon usage is considered among the important features and used as part of the feature set in synonymous effect prediction (Buske OJ et al, 2013; Livingstone M et al 2017; Zhang X et al 2017). It was however, observed that the RSCU of alternate allele and the difference with the reference is used for the prediction and the reference value is not used by any predictor. In addition to global RSCU values, transcript based RSCU values were also found to be significant.

In addition to splice regulation disruption annotation that is used in synonymous effect prediction, we used the ESR score of the reference, alternate allele and their difference. A similar pattern was observed for this feature group with ESR score of the reference allele being reported as important in addition to the ESR score of the alternate and the difference between the reference and alternate.

The identity of the 2^nd^ and 3^rd^ bases in the reference codon was significant in several of the models. We speculate that these could have direct correlation to the conservation of that position.

Apart from the above features, the length and distance of the variant from the exon end also contributed to the model prediction.

Our results suggest that there can be inherent significance in the reference that is independent of the alternate allele. Feature selection for some genomic properties might thus benefit from such expansion of the underlying features and the inclusion of reference features in model building. We started with a basic set of commonly used features and the approach can be expanded to other features for evaluation.

A class segregation that more closely mimics reality would be an additional transitional class with moderate conservation in addition to high conservation and low conservation classes. Our models built around this performed almost at par and features underlying the model remained similar. Variant predictors could also extend the models to cover variants in the first and last exons and single exon genes which might potentially show different behaviour based on their location and require separate models with a corresponding set of features. Indeed, other predictors (Livingstone

M et al, 2017), have observed differences in the significant features and prediction performance based on the distance of variants along the exon and proposed different models to address this.

Features selected from our model showed a reasonable replacement for conservation in a synonymous effect prediction tool and showed potential for further improvement of prediction when added to the orginal features of the tool.

## Conclusion

Our novel approach helped identify key features that can be used to build better models for synonymous mutations. Moreover, the identification of significant reference features that are independent of alternate alleles provides interesting avenues for further exploration to identify vulnerable sites in the genome and also gain insight into possible disease mechanisms.

## Supplementary Tables and Figures

**Supplementary Fig 1:**
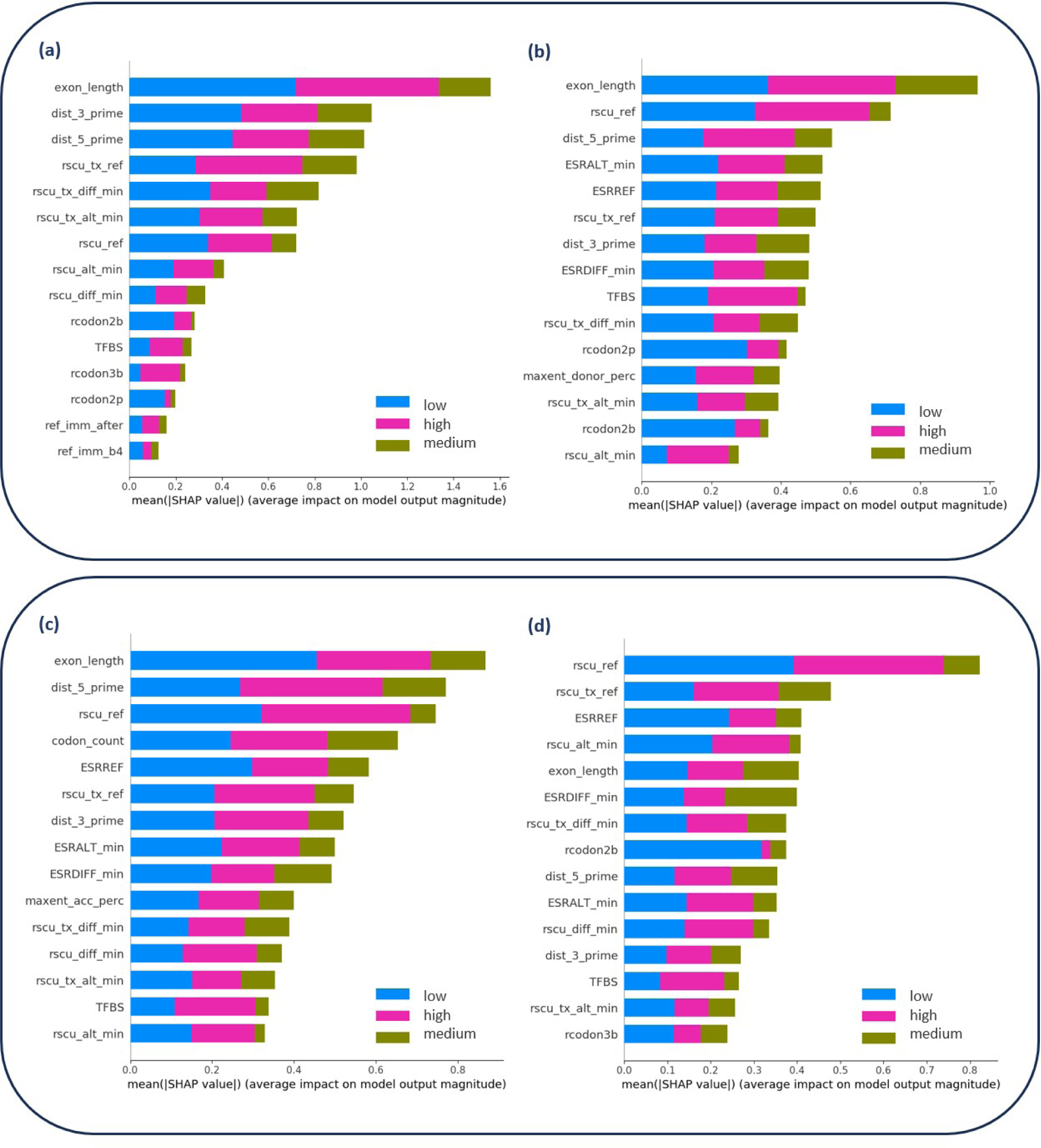
Significant features based on mean SHAP values for (clockwise) single exon, first exon, last exon and general models with high class PP100 >= 90 percentile

**Supplementary Table 1:**
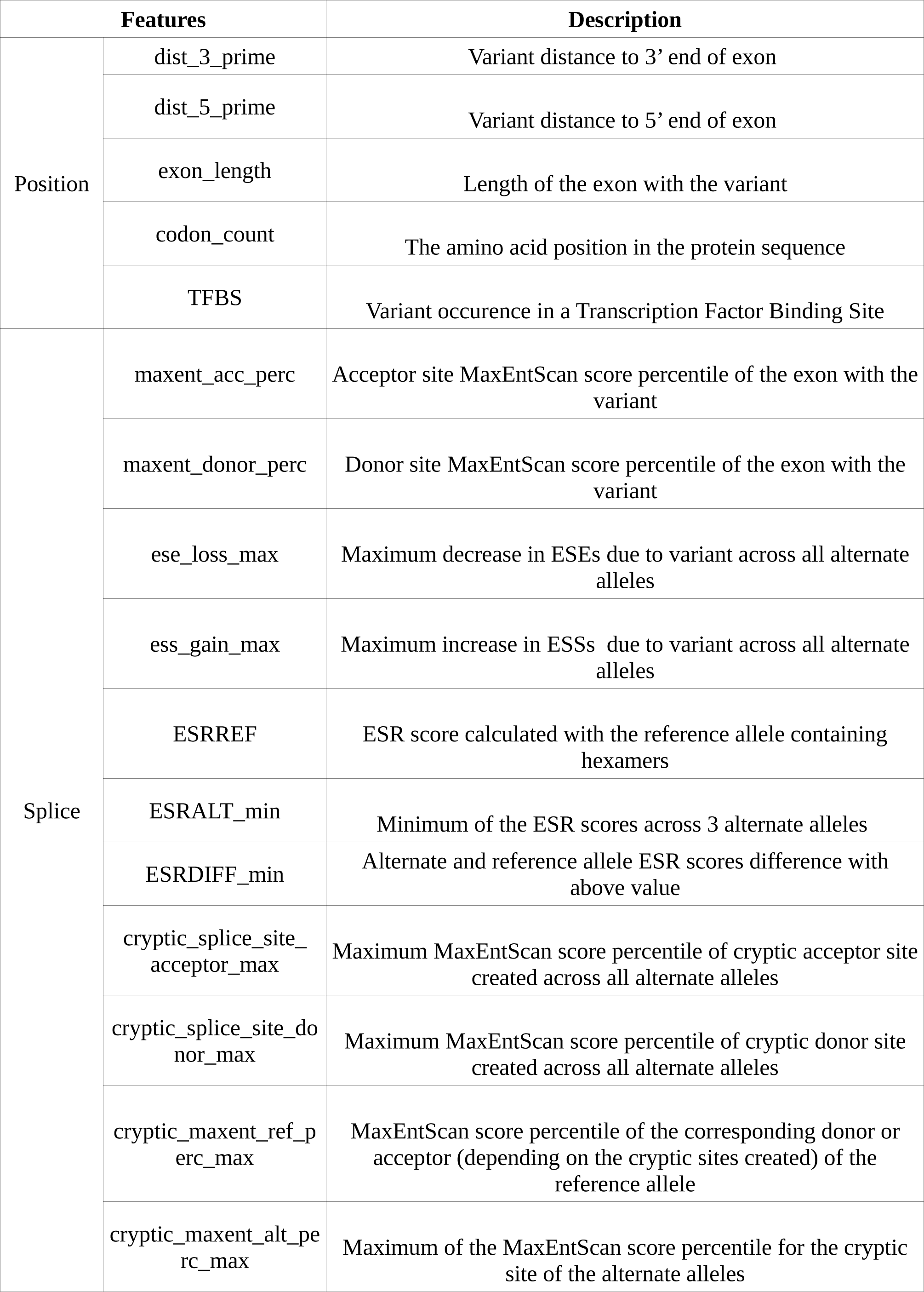
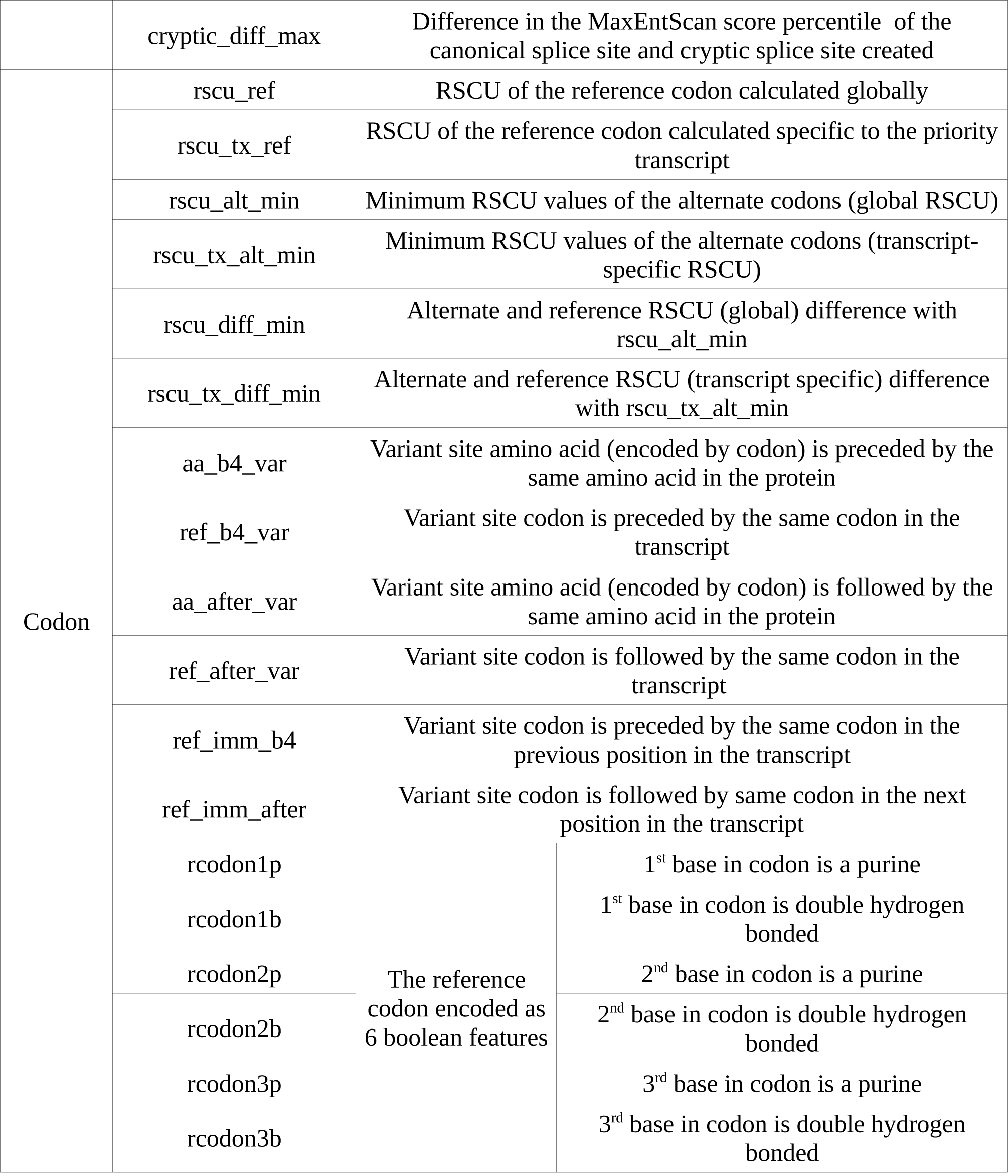
Features used to build the 3 class models.

| Features |  | Description |
| --- | --- | --- |
| Position | dist_3_prime | Variant distance to 3' end of exon |
|  | dist_5_prime | Variant distance to 5' end of exon |
|  | exon_length | Length of the exon with the variant |
|  | codon_count | The amino acid position in the protein sequence |
|  | TFBS | Variant occurrence in a Transcription Factor Binding Site |
| Splice | maxent_acc_perc | Acceptor site MaxEntScan score percentile of the exon with the variant |
|  | maxent_donor_perc | Donor site MaxEntScan score percentile of the exon with the variant |
|  | ese_loss_max | Maximum decrease in ESEs due to variant across all alternate alleles |
|  | ess_gain_max | Maximum increase in ESSs due to variant across all alternate alleles |
|  | ESRREF | ESR score calculated with the reference allele containing hexamers |
|  | ESRALT_min | Minimum of the ESR scores across 3 alternate alleles |
|  | ESRDIFF_min | Alternate and reference allele ESR scores difference with above value |
|  | cryptic_splice_site_acceptor_max | Maximum MaxEntScan score percentile of cryptic acceptor site created across all alternate alleles |
|  | cryptic_splice_site_donor_max | Maximum MaxEntScan score percentile of cryptic donor site created across all alternate alleles |
|  | cryptic_maxent_ref_perc_max | MaxEntScan score percentile of the corresponding donor or acceptor (depending on the cryptic sites created) of the reference allele |
|  | cryptic_maxent_alt_perc_max | Maximum of the MaxEntScan score percentile for the cryptic site of the alternate alleles |

**Supplementary Table 1:** Features used to build the 3 class models
|  |  |  |  |
| --- | --- | --- | --- |
|  | cryptic_diff_max | Difference in the MaxEntScan score percentile of the canonical splice site and cryptic splice site created |  |
| Codon | rscu_ref | RSCU of the reference codon calculated globally |  |
|  | rscu_tx_ref | RSCU of the reference codon calculated specific to the priority transcript |  |
|  | rscu_alt_min | Minimum RSCU values of the alternate codons (global RSCU) |  |
|  | rscu_tx_alt_min | Minimum RSCU values of the alternate codons (transcript-specific RSCU) |  |
|  | rscu_diff_min | Alternate and reference RSCU (global) difference with rscu_alt_min |  |
|  | rscu_tx_diff_min | Alternate and reference RSCU (transcript specific) difference with rscu_tx_alt_min |  |
|  | aa_b4_var | Variant site amino acid (encoded by codon) is preceded by the same amino acid in the protein |  |
|  | ref_b4_var | Variant site codon is preceded by the same codon in the transcript |  |
|  | aa_after_var | Variant site amino acid (encoded by codon) is followed by the same amino acid in the protein |  |
|  | ref_after_var | Variant site codon is followed by the same codon in the transcript |  |
|  | ref_imm_b4 | Variant site codon is preceded by the same codon in the previous position in the transcript |  |
|  | ref_imm_after | Variant site codon is followed by same codon in the next position in the transcript |  |
|  | rcodon1p | The reference codon encoded as 6 boolean features | 1<sup>st</sup> base in codon is a purine |
|  | rcodon1b |  | 1<sup>st</sup> base in codon is double hydrogen bonded |
|  | rcodon2p |  | 2<sup>nd</sup> base in codon is a purine |
|  | rcodon2b |  | 2<sup>nd</sup> base in codon is double hydrogen bonded |
|  | rcodon3p |  | 3<sup>rd</sup> base in codon is a purine |
|  | rcodon3b |  | 3<sup>rd</sup> base in codon is double hydrogen bonded |

**Supplementary Table 2:**
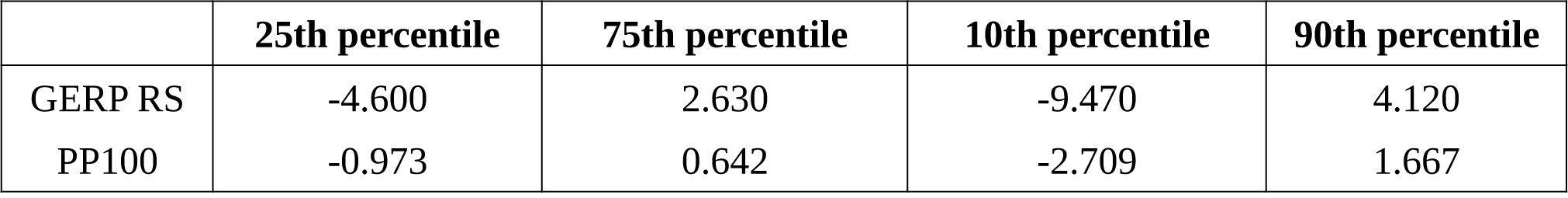
Class cutoff values for GERP and PhyloP models.

**Supplementary Table 3:**
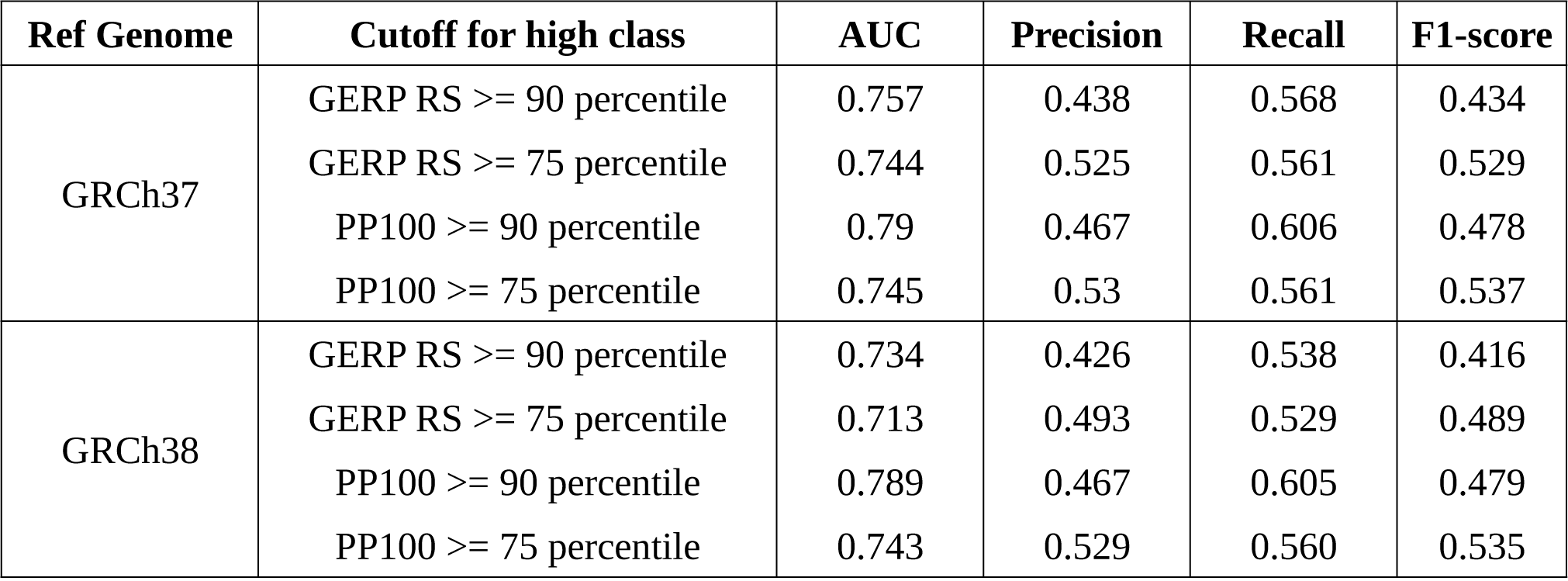
GRCh37 vs GRCh38 results from independent test set.

**Supplementary Table 4:** Individual feature models performance with high class PhyloP100 >= 90 percentile of dataset.

| Feature | AUC | Precision | Recall | F1-score |
| --- | --- | --- | --- | --- |
| rscu_ref | 0.727 | 0.412 | 0.528 | 0.361 |
| rscu_diff_min | 0.727 | 0.412 | 0.528 | 0.361 |
| rscu_alt_min | 0.703 | 0.400 | 0.511 | 0.330 |
| rscu_tx_ref | 0.642 | 0.380 | 0.458 | 0.314 |
| rscu_tx_diff_min | 0.634 | 0.376 | 0.450 | 0.311 |
| ESRREF | 0.617 | 0.373 | 0.432 | 0.290 |
| ESRALT_min | 0.611 | 0.371 | 0.437 | 0.279 |
| rcodon2b | 0.604 | 0.099 | 0.470 | 0.159 |
| rscu_tx_alt_min | 0.588 | 0.357 | 0.405 | 0.221 |
| ESRDIFF_min | 0.575 | 0.372 | 0.406 | 0.328 |
| exon_length | 0.565 | 0.357 | 0.384 | 0.230 |
| rcodon3b | 0.542 | 0.326 | 0.379 | 0.242 |
| ese_loss_max | 0.533 | 0.078 | 0.373 | 0.118 |
| TFBS | 0.530 | 0.313 | 0.364 | 0.319 |
| rcodon2p | 0.528 | 0.322 | 0.365 | 0.168 |

**Supplementary Table 4:** Individual feature models performance with high class PhyloP100 $\geq 90$ percentile of dataset.
| **Feature** | **AUC** | **Precision** | **Recall** | **F1-score** |
| --- | --- | --- | --- | --- |
| dist_5_prime | 0.523 | 0.343 | 0.351 | 0.189 |
| rcodon3p | 0.523 | 0.312 | 0.361 | 0.277 |
| ess_gain_max | 0.519 | 0.073 | 0.356 | 0.110 |
| dist_3_prime | 0.518 | 0.343 | 0.348 | 0.194 |
| codon_count | 0.511 | 0.336 | 0.341 | 0.227 |
| rscu_group | 0.750 | 0.429 | 0.562 | 0.401 |
| splice group | 0.682 | 0.409 | 0.500 | 0.390 |
| rcodon_group | 0.718 | 0.405 | 0.524 | 0.318 |
| All 3 groups together | 0.785 | 0.460 | 0.604 | 0.465 |
| Top 10 feat (from single feat models) | 0.778 | 0.453 | 0.597 | 0.453 |
| Top 20 | 0.789 | 0.464 | 0.607 | 0.473 |

**Supplementary Table 5:**
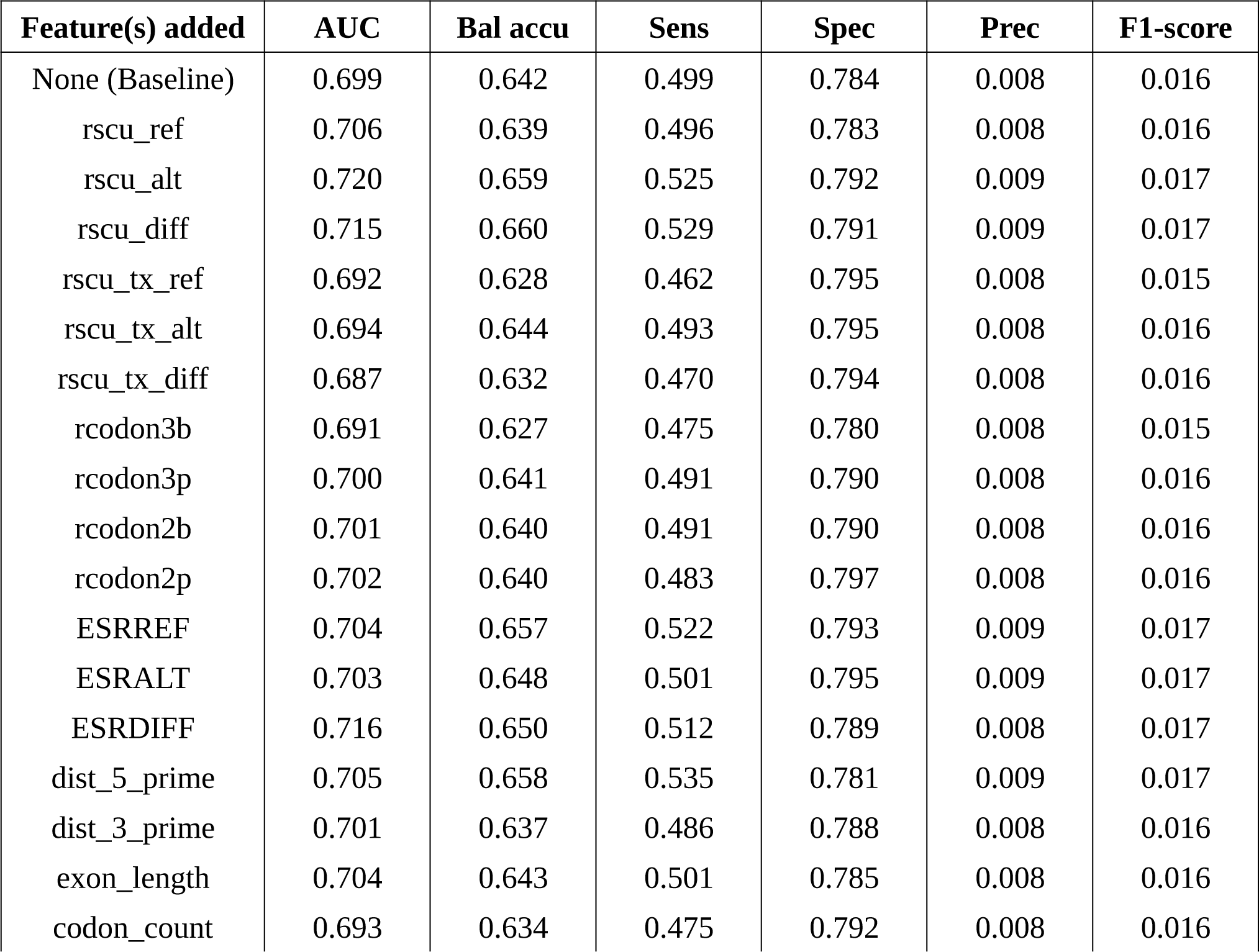
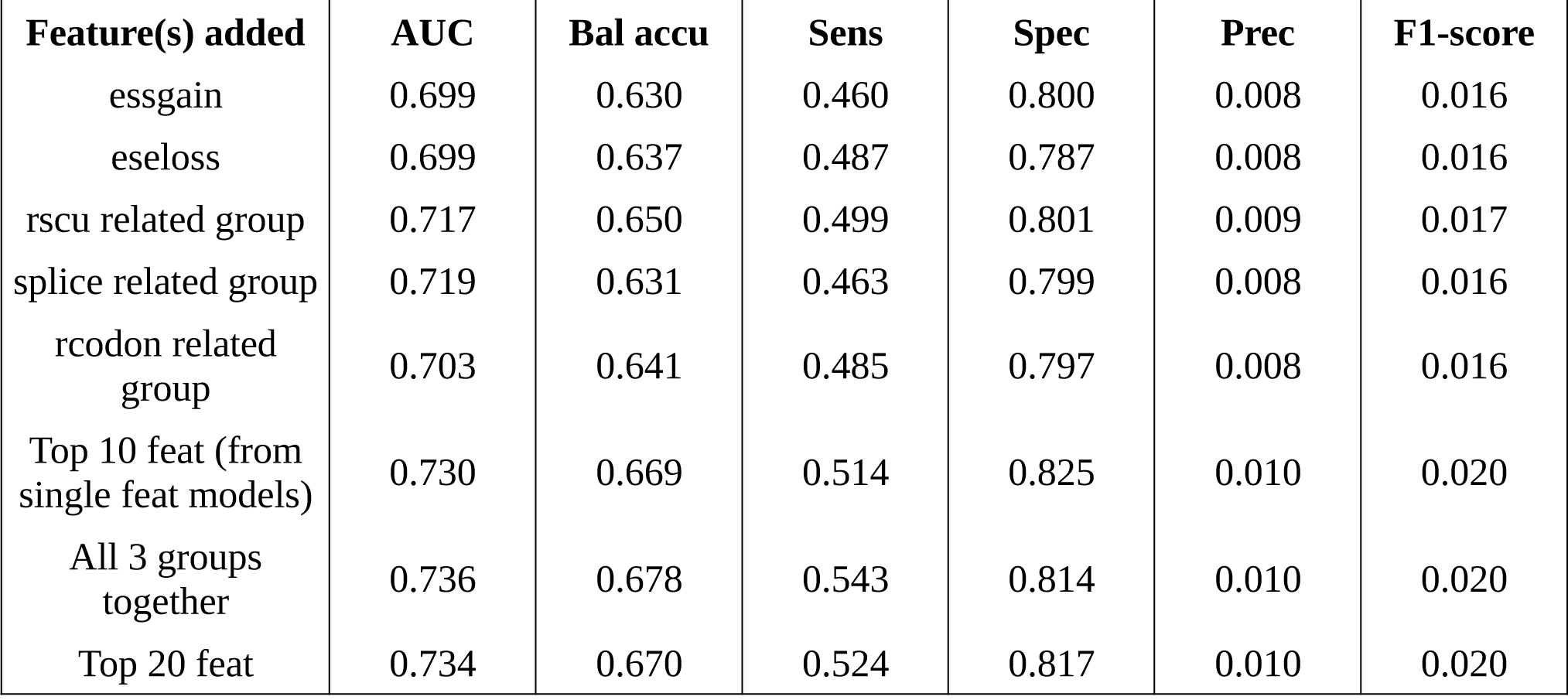
Test set results with IDSV features without conservation (PhyloP) feature with individual features and feature groups added (average reults over 10 runs)

